# Yomix: An Interactive Tool for the Exploration of Low-Dimensional Embeddings in Omics Data

**DOI:** 10.64898/2025.12.05.692343

**Authors:** Nicolas Perrin-Gilbert, Nisma Amjad, Pierre Fumeron, Silvia Tulli, Joshua J. Waterfall

**Affiliations:** Sorbonne Université, CNRS, Institut des Systèmes Intelligents et de Robotique, ISIR, F-75005, Paris, France; Inserm U1330, Institut Curie Centre de Recherche, PSL University, Paris, France; Translational Research Department, Institut Curie Centre de Recherche, PSL University, Paris, France

**Author notes:** Correspondence email addresses.

## Abstract

**Summary:** In the analysis of diverse omics data, a common and important preliminary step involves computing low-dimensional embeddings using techniques such as PCA, UMAP, t-SNE, or variational autoencoders. These embeddings provide a global overview of sample distributions and their relationships, often serving as the basis for formulating biological hypotheses. To facilitate rapid and intuitive exploration of such low-dimensional embeddings, we developed Yomix, an interactive omics-agnostic visualization and data exploration tool. Yomix enables users to flexibly define subsets of interest using a lasso selection tool, instantly compute their feature signatures, and compare their distributions. Yomix is a fast and efficient tool for interactive exploration of diverse omics datasets.

**Availability and Implementation:** Yomix and its documentation are publicly available at https://github.com/perrin-isir/yomix.

## 1 Introduction

The high-dimensional nature of omics datasets presents substantial challenges in data interpretation. To facilitate visualization and exploration of such datasets, a widely adopted preliminary step is the computation of low-dimensional embeddings, typically in two or three dimensions (Becht et al., 2018; Moon et al., 2019). Dimensionality reduction techniques, such as Principal Component analysis (PCA), Uniform Manifold Approximation and Projection (UMAP) (McInnes et al., 2018), t-distributed Stochastic Neighbour Embedding (t-SNE) (van der Maaten, 2009), and variational autoencoders (VAEs) (Lopez et al., 2018), are frequently employed for this purpose, offering advantages in capturing both global and local data features. These low-dimensional representations serve as a basis for deriving biological insights, enabling researchers to identify clusters (Kiselev et al., 2019), infer developmental trajectories (Saelens et al., 2019), and formulate hypotheses regarding underlying biological processes (Trapnell et al., 2014).

Despite the widespread adoption of dimensionality reduction techniques in omics workflows, effectively interpreting the resulting low-dimensional embeddings remains a significant challenge. In practice, researchers must frequently explore these embeddings iteratively, zooming into local regions, selecting subpopulations of interest, and querying their molecular signatures. This type of visual exploration is essential for identifying novel cell types, validating marker features (genes, CpG sites, k-mers, etc.). However, most existing analysis frameworks (e.g., Seurat (Hao et al., 2021), Scanpy (Wolf et al., 2018)) rely on static visualizations that are not directly responsive to user interaction or dynamic filtering. This restricts the user’s ability to adjust views, subset groups, or overlay metadata in real time, often requiring time-consuming re-computation or manual scripting for even basic exploratory tasks. While recent tools have been developed to address some of these issues (Kotliar and Colubri, 2021; Keller et al., 2025; Li et al., 2020) there remains a significant need for fast, flexible approaches to interact with diverse types of omics datasets..

To address these challenges, we developed Yomix, a lightweight yet powerful visualization framework that supports interactive projection views, metadata integration, and feature ranking across a variety of omics data types.

## 2 Materials and Methods

Yomix is a lightweight tool used to identify biologically relevant structures and discriminative features across multiple datasets. It is developed in Python and uses the Bokeh framework, which generates a JavaScript-based graphical interface and callbacks that run in the web browser. It requires an anndata object (Virshup et al., 2024) saved in.h5ad format as input which contains a data matrix and at least one low-dimensional embedding.

By launching Yomix from the Python session, it directly opens an interactive plot of the embedding chosen by the user (Figure 1A-E). Users can switch between different embeddings, change plot sizes and labels, and define specific subsets (Figure 1A). These subsets can be selected using the legend and predefined annotations (Figure 1B) or manually with a lasso tool (Figure 1C), making it highly interactive. Yomix is both omics agnostic and handles three-dimensional (3D) embeddings. It also supports embeddings with more than 3 dimensions by letting users pick amongst dimensions for every axis.

**Figure 1:**
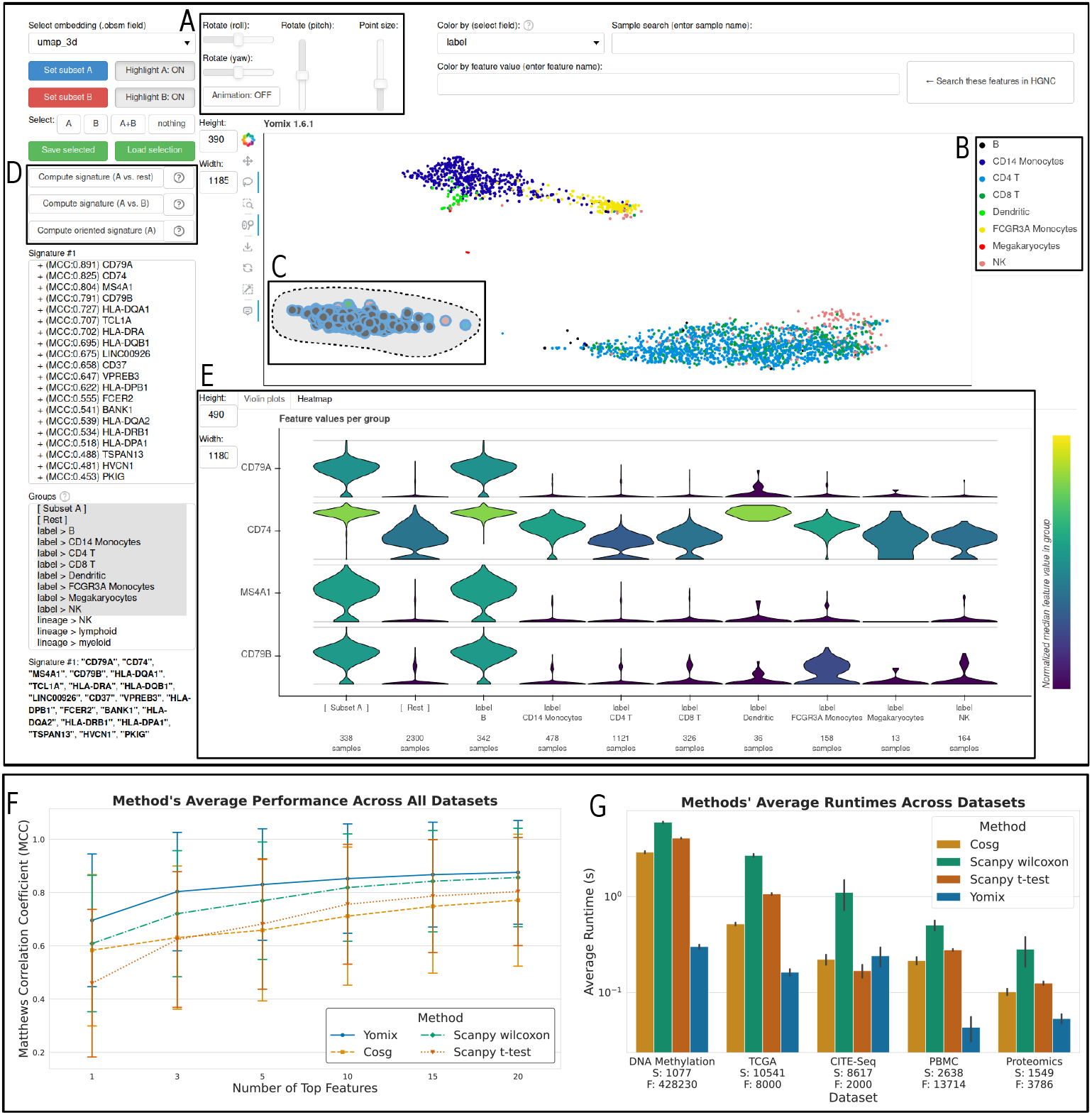
Screenshot of Yomix’s interface displaying the PBMC dataset after computing the signature of the bottom-left cluster, mostly composed of B cells. A) Buttons to control real-time 3D rotation, and to interact in 3D with the scatter plot. B) Users can click on legend to select data. C) Lasso tool for manual selection of data points. D) Users can compute different types of signatures. E) Violin plots to visualize the distribution of selected features for selected subgroups. Users can switch to a heat map format. F) Average performance comparison between SVM trained on features selected by different methods, error bars represent standard deviation. G) Runtime comparison between the different methods, error bars represent standard deviation

Yomix can identify signatures for a user-defined cluster against the remaining data or between two user-defined subsets (Figure 1D). These signatures, or marker features, are computed using a rapid, two-step filtering process that identifies the most discriminative features.

The algorithm first computes group statistics for each feature, calculating mean and standard deviation for Group A and Group B (either a specific subset or the remaining samples). It then calculates the Wasserstein distance between distributions for each feature, ranks all features in descending order by Wasserstein distance, and selects the features with the highest distributional differences. The second step identifies the features among the pre-filtered set that most reliably distinguish between groups using optimal binary classification thresholds. Specifically, for each feature, the algorithm tests multiple thresholds across the range of feature values, computes the confusion matrix for each threshold, calculates the Matthews Correlation Coefficient (MCC) at each threshold, and selects the one that maximizes MCC. The computation is vectorized across all features and thresholds simultaneously for efficient computation. A key advantage of this approach is its speed and accuracy, as filtering reduces the search space, and vectorized operations accelerate the search of optimal thresholds. More details on this implementation are available in Yomix documentation https://perrin-isir.github.io/yomix/

In addition, Yomix can compute oriented signatures by letting the user select a subset and draw an arrow in the embedding space. Each element of the subset is projected onto the arrow, creating a vector of positions along the arrow. For every feature, the algorithm computes the Pearson correlation between the feature values and the projection vector. It then returns the top features sorted by highest correlation. Selected features, either from signatures or input by the user, are visualized separately as either violin plots or heatmaps (Figure 1E). Analyses were run on a Dell desktop computer equipped with 32.0 GiB of RAM and a 13th Gen Intel® Core™ i9-13950HX processor.

## 3 Results

To evaluate the predictive and computational performance of Yomix, we compared it with three widely used feature selection methods: *t* test, Wilcoxon test, and COSG, which was shown to perform well in a recent benchmarking study (Pullin and McCarthy, 2024). Benchmarks were performed on five diverse omics datasets that include bulk RNAseq from The Cancer Genome Atlas (TCGA) pancancer analysis (Hoadley, 2018), DNA methylation array profiling of over 1000 sarcomas (Koelsche et al., 2021), single cell multiomics of cord blood (Stoeckius et al., 2017), scRNAseq of peripheral blood mononuclear cells (PBMCs) (Hansen et al., 2023) and proteomics from the National Cancer Institute’s Clinical Proteomic Tumor Analysis Consortium (CPTAC) (Li et al., 2023).

To evaluate the utility of features selected by the various methods, we selected an increasing number of top features (from 1 to 20) from each method and measured the average MCC using a support vector machine (SVM) as the classifier with default parameters on these features across all datasets. We assessed feature signature quality using MCC scores from the SVM classification.

The results shown in the Figure 1 demonstrate that Yomix consistently selects feature signatures with the highest predictive power (Figure 1F). Yomix achieved average MCC as good as that of other methods across nearly all feature set sizes, underscoring its ability to select discriminative and biologically informative markers. Wilcoxon was the second best performing method, while COSG and *t* test yielded feature sets with lower MCC scores. All methods showed improved performance when more features were included.

In addition to predictive performance, speed was also computed for each method. The shortest runtimes in all datasets were also attained by Yomix, with execution times remaining below 0.7 seconds even for the largest datasets (Figure 1G). When analyzing the TCGA bulk RNAseq dataset, which has 10,541 samples and 8,000 features, the signature computation was completed in approximately 0.165 seconds. In contrast, the Wilcoxon method exhibited the longest runtime, 5.89 seconds on methylation and 2.99 seconds on TCGA datasets. COSG and *t* test methods displayed intermediate performance as they outperformed Wilcoxon but remained slower than Yomix across all datasets. The results demonstrate the significant computational efficiency of Yomix. These findings highlight the scalability and speed of Yomix’s two-step algorithm, making it particularly well-suited for the rapid, iterative analysis required in large-scale omics exploration.

This analysis confirms that Yomix not only offers superior computational speed and interactivity but also excels at identifying feature sets with notable biological relevance and predictive capability, regardless of the type of dataset used and the size of samples required.

## Funding

This work was supported by ITMO Cancer of Aviesan within the framework of the 2021–2030 Cancer Control Strategy, on funds administered by Inserm [22CM060-00].

## Notes

### Competing Interest Statement

The authors have declared no competing interest.

